## Supplementary material for "Modulation of fatty acid elongation sustains sexually dimorphic hydrocarbons and female attractiveness in *Blattella germanica* (L.)": S1 Appendix

Multiple alignment of BgElo protein sequences, homologous fragments are highlighted in light blue or burgundy, conserved motifs are labeled, including a HXXHH motif and a YXYY motif. Some BgElos missed the motifs due to incomplete cloning of the CDS.

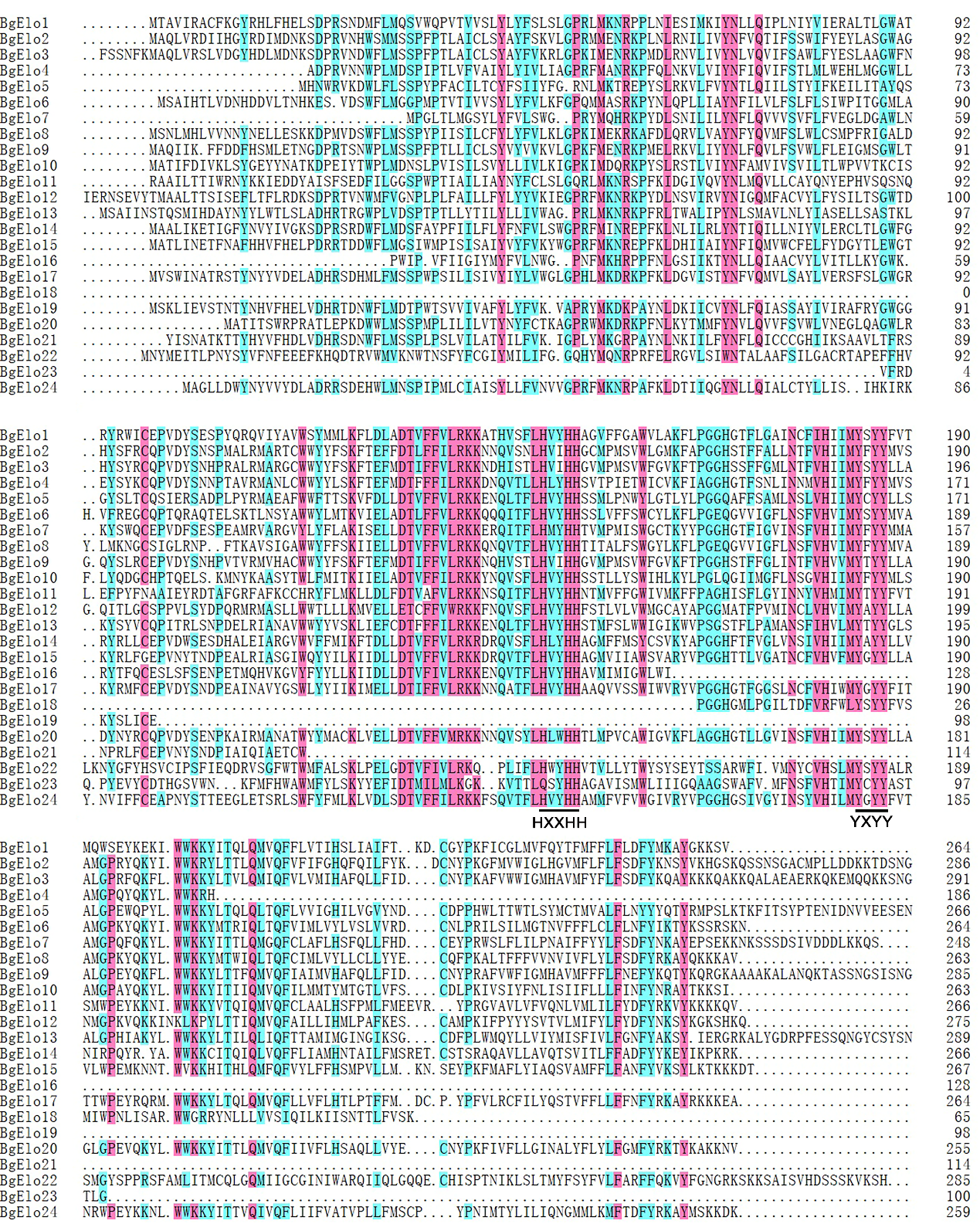

Gene location, intron-exon structure, protein structure of different BgElos. Different numbers above the introns or under the exons represent the length of nucleic acid sequences, the numbers above the unknown region or Elo domain represent the number of amino acid residues. Locations of the first exon in Scaffold are marked by black numbers (the position of the first nucleic acid base) behind the Scaffold and black vertical lines.

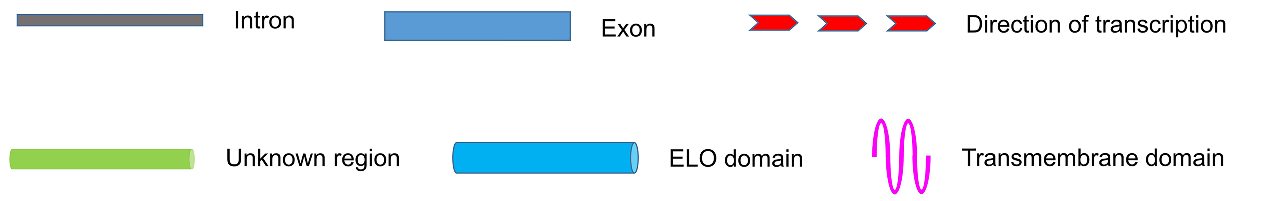

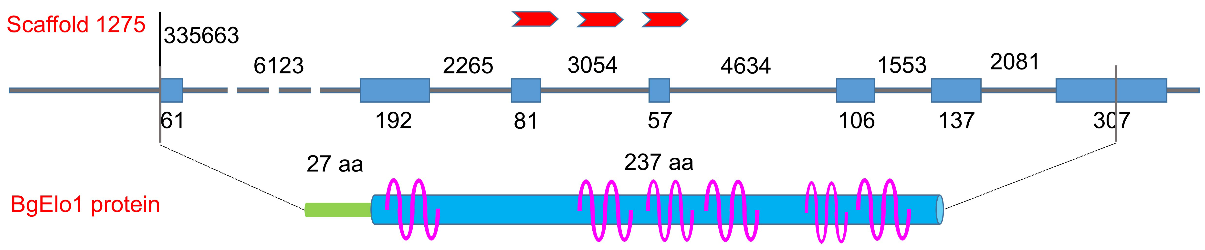

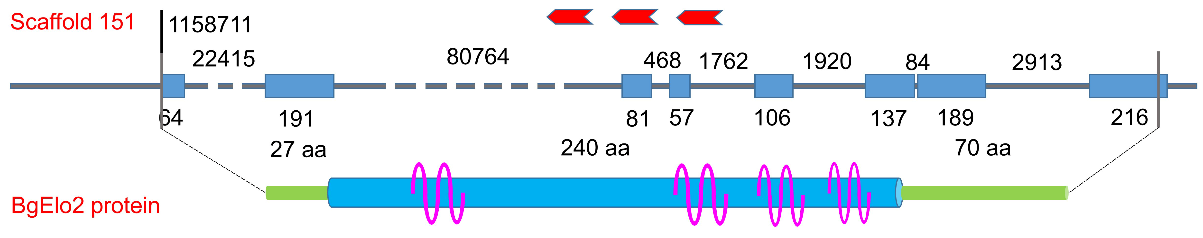

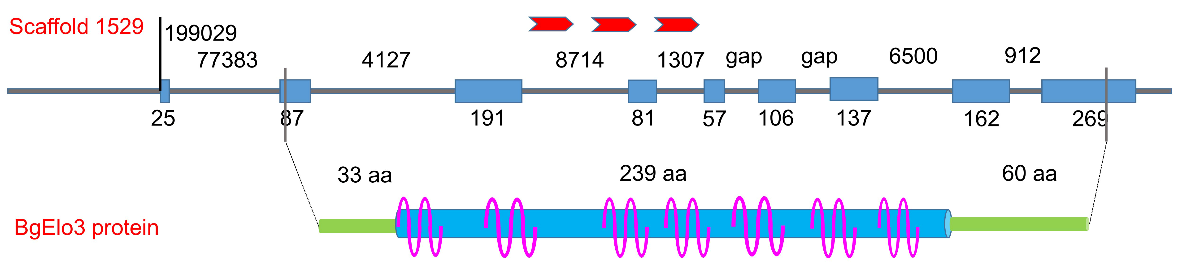

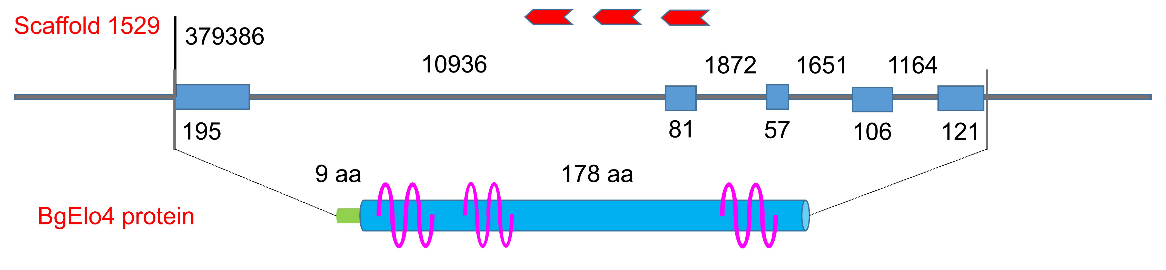

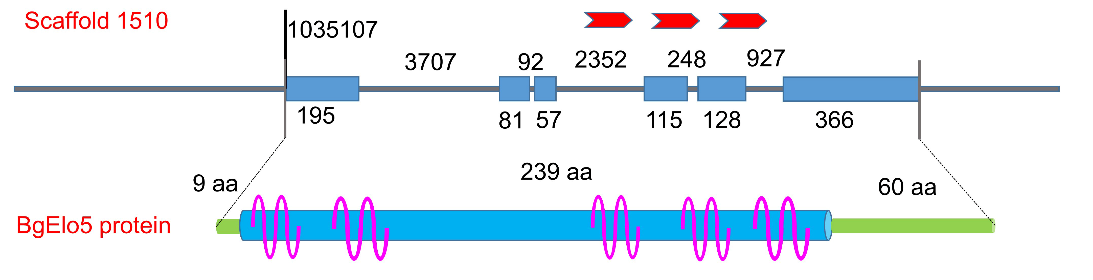

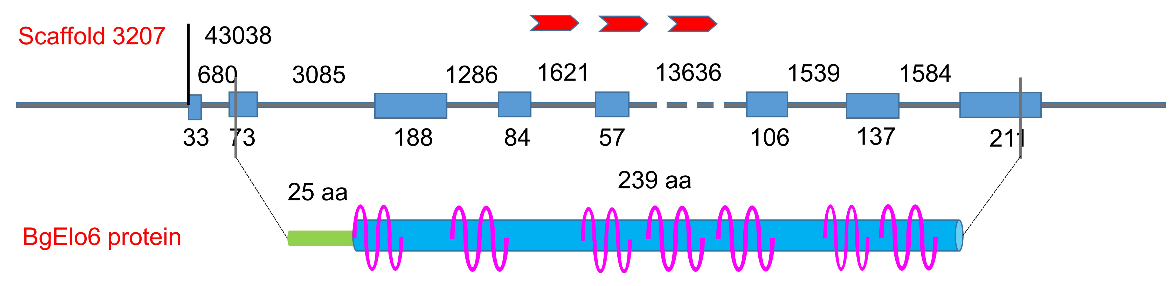

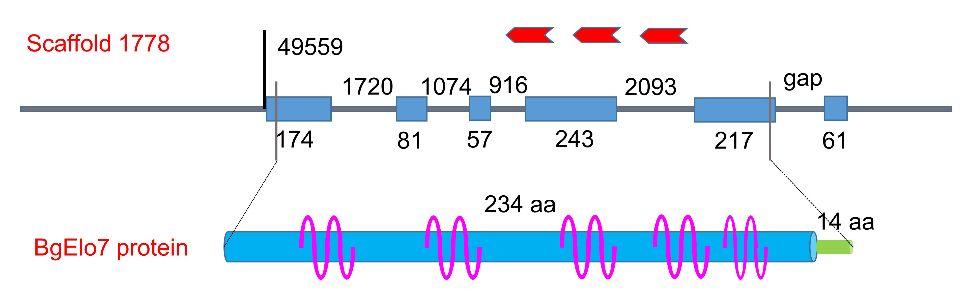

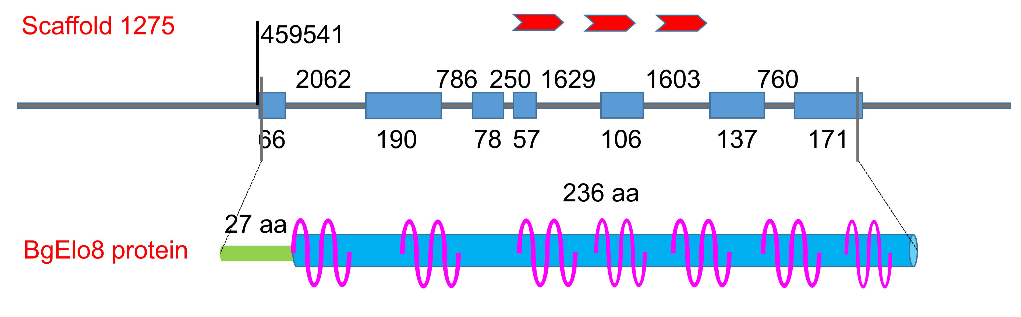

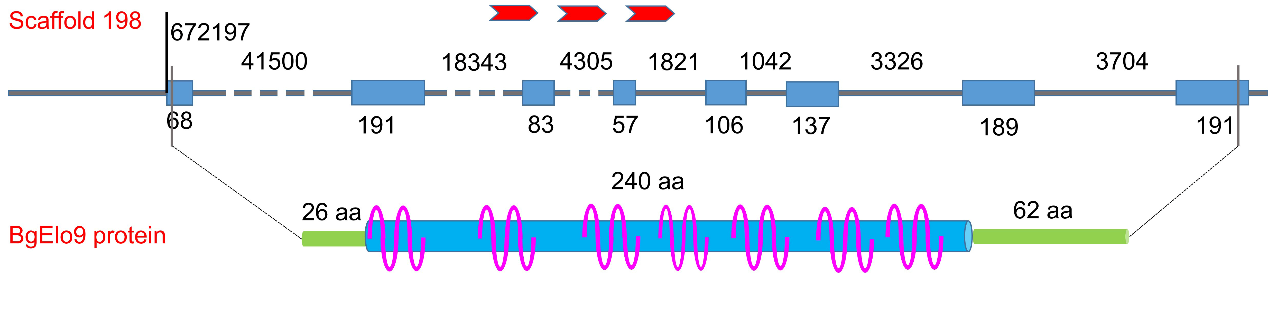

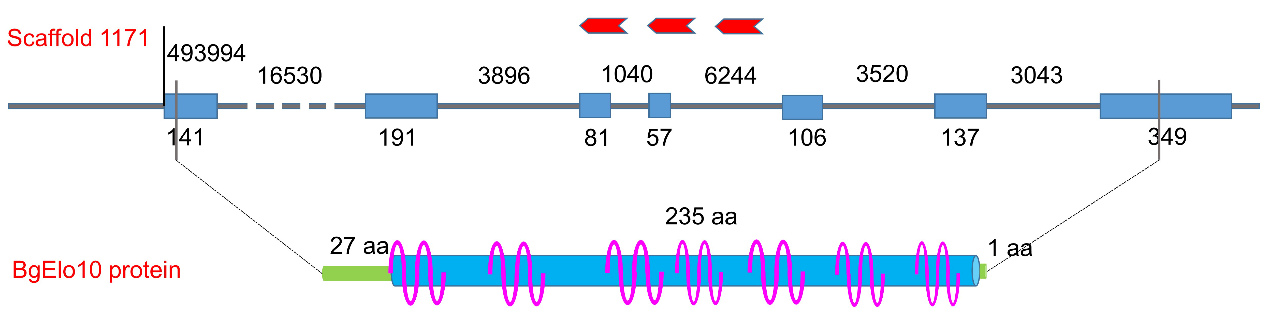

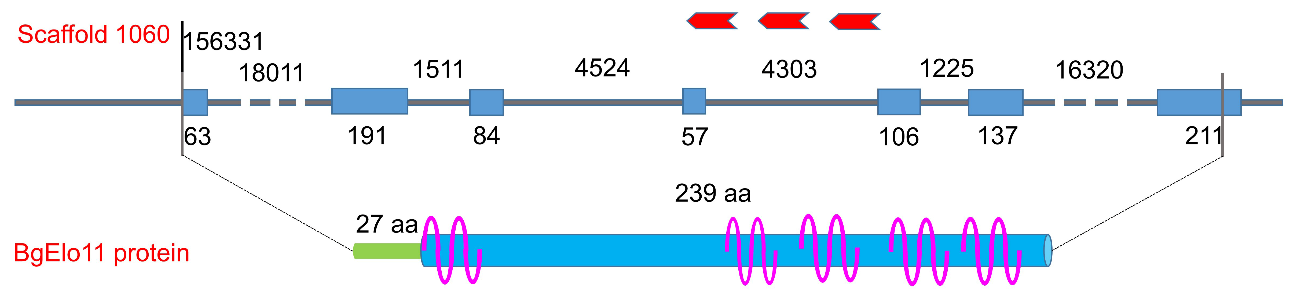

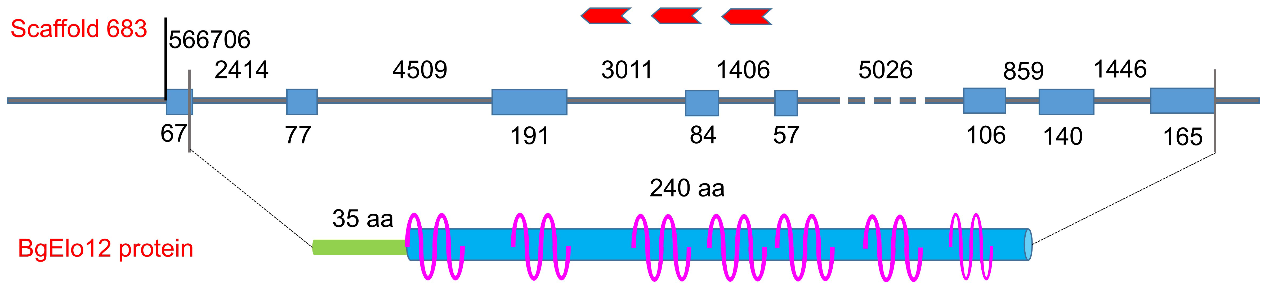

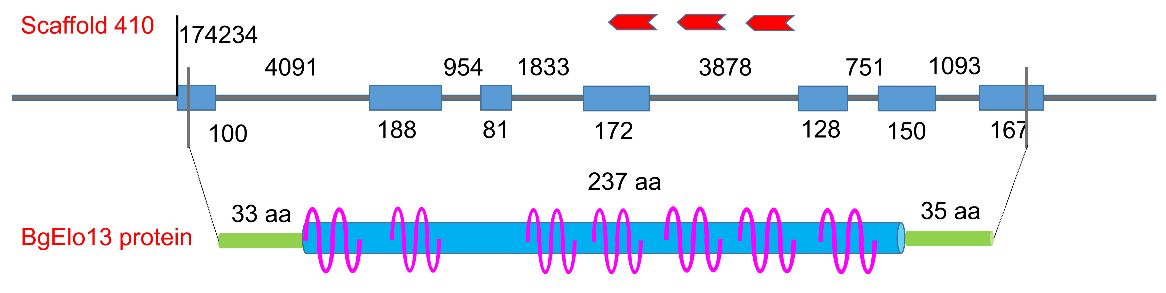

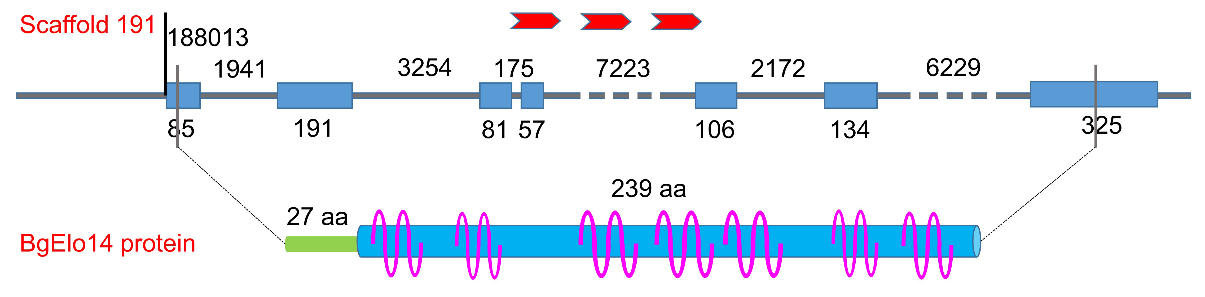

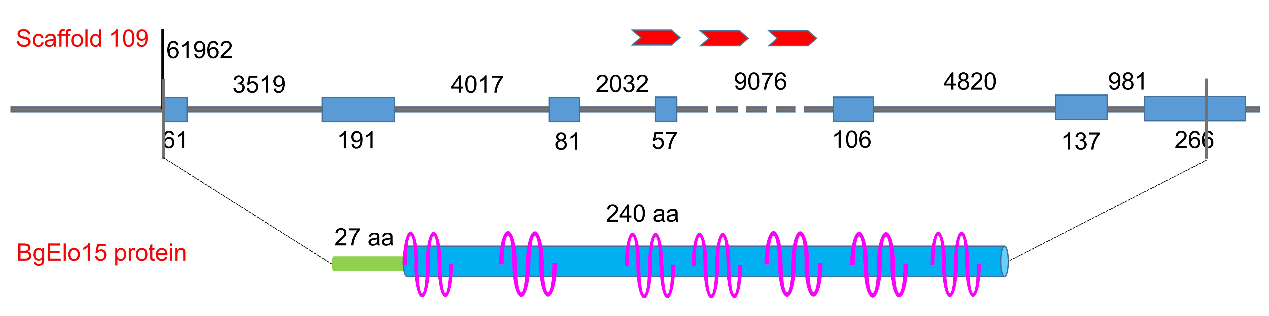

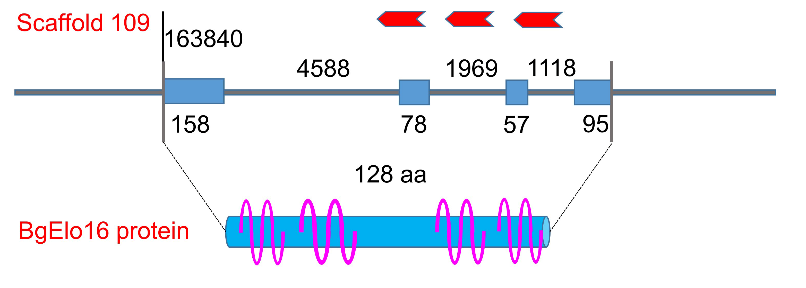

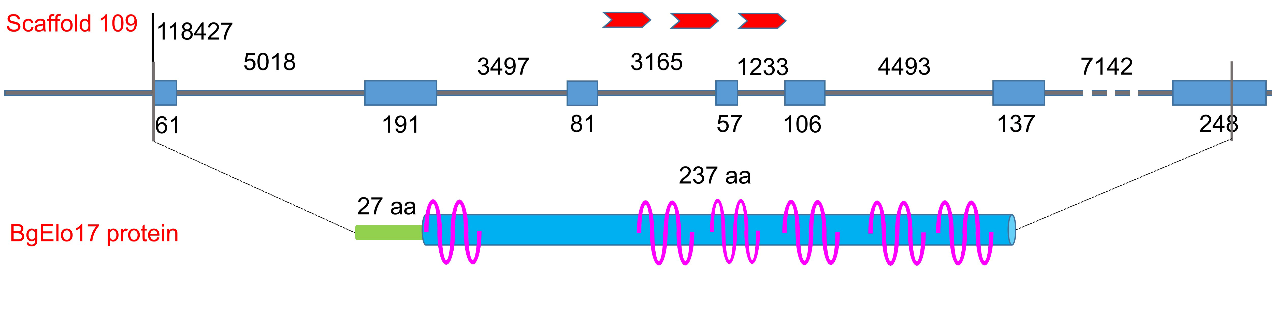

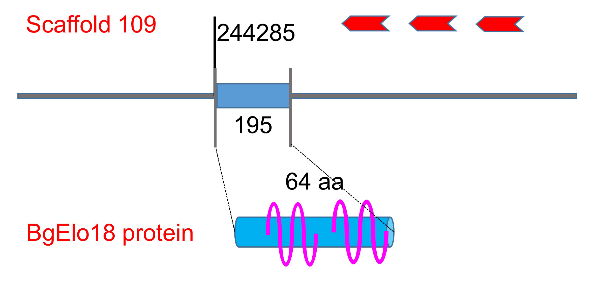

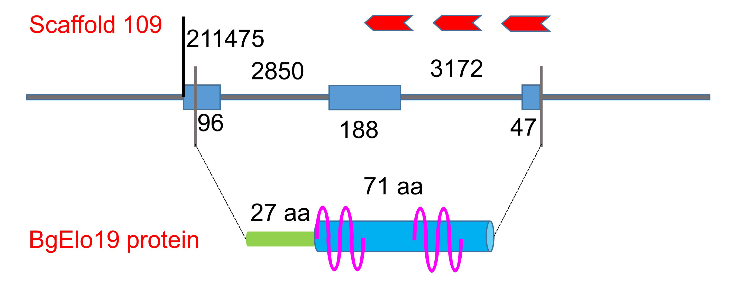

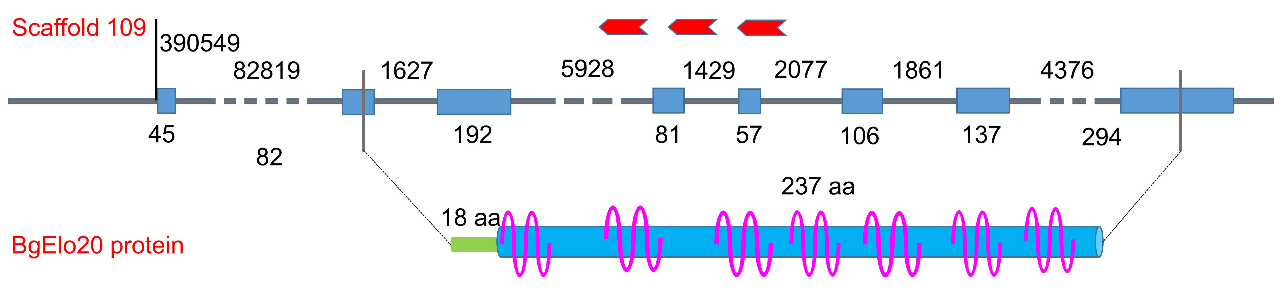

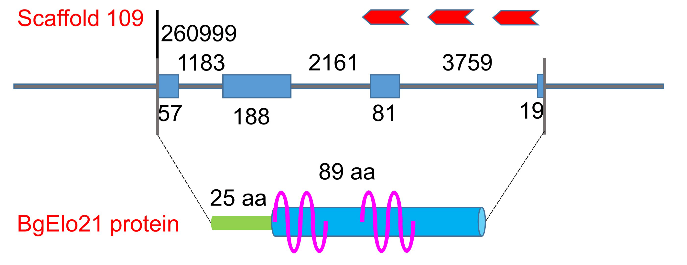

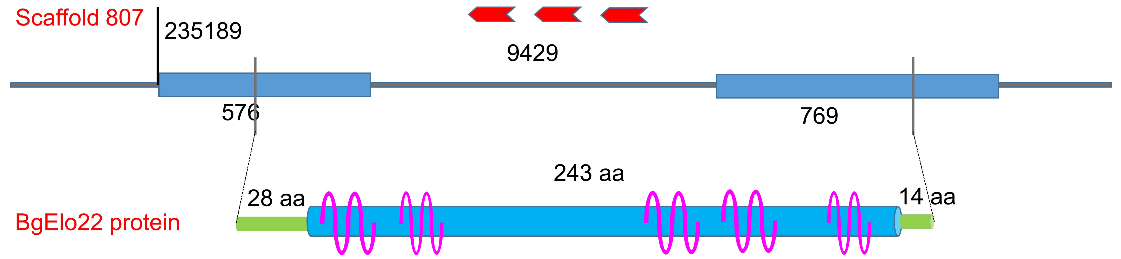

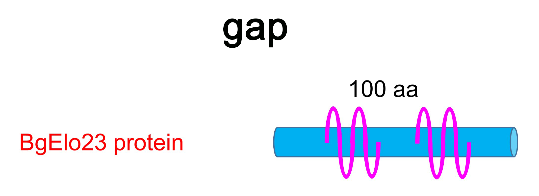

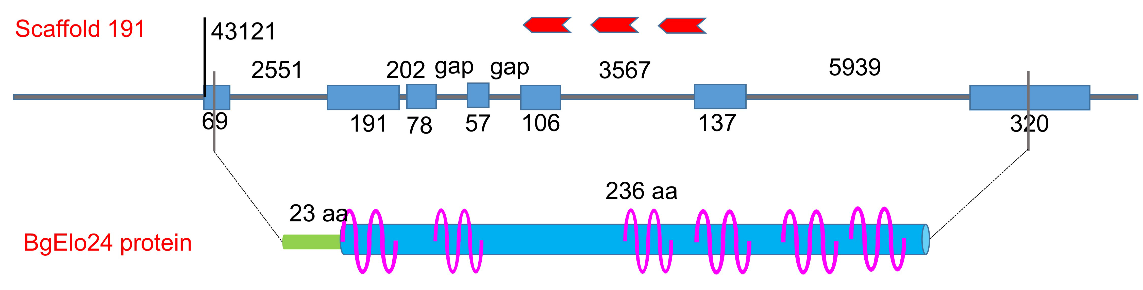
