## Supplementary material for "Modulation of fatty acid elongation sustains sexually dimorphic hydrocarbons and female attractiveness in *Blattella germanica* (L.)": S1Table

| **Quantification of individual cuticular hydrocarbon from AD1–AD6 female and male cockroaches.** | | | | | | | | |
| --- | --- | --- | --- | --- | --- | --- | --- | --- |
| **Peak No.** | **Compound** | **AD1-female** | | **AD2-female** | **AD3-female** | **AD4-female** | **AD5-female** | **AD6-female** |
| **1** | **n-C27** | 1.326 ± 0.183 | | 1.220 ± 0.111 | 1.170 ± 0.277 | 1.423 ± 0.137 | 1.162 ± 0.126 | 1.484 ± 0.174 |
| **2** | **11-; 13-MeC27** | 1.303 ± 0.196 | | 1.319 ± 0.172 | 1.260 ± 0.245 | 1.505 ± 0.130 | 1.487 ± 0.175 | 1.461 ± 0.185 |
| **3** | **5-MeC27** | 1.522 ± 0.221 | | 1.537 ± 0.194 | 1.365 ± 0.293 | 1.562 ± 0.124 | 1.383 ± 0.161 | 1.454 ± 0.175 |
| **4** | **11,15-DimeC27** | 0.199 ± 0.036 | | 0.213 ± 0.026 | 0.213 ± 0.040 | 0.427 ± 0.067 | 0.319 ± 0.040 | 0.334 ± 0.044 |
| **5** | **3-MeC27** | 1.985 ± 0.283 | | 2.010 ± 0.229 | 1.876 ± 0.341 | 2.026 ± 0.146 | 2.166 ± 0.231 | 2.393 ± 0.258 |
| **6** | **5, 9-; 5,11-DimeC27** | 0.852 ± 0.085 | | 0.976 ± 0.142 | 1.273 ± 0.125 | 1.643 ± 0.161 | 1.086 ± 0.178 | 0.914 ± 0.143 |
| **7** | **n-C28** | 0.834 ± 0.041 | | 0.904 ± 0.038 | 0.855 ±0.055 | 0.821 ± 0.062 | 0.746 ± 0.030 | 0.865 ± 0.049 |
| **8** | **3, 9-; 3,11-DimeC27** | 0.860 ± 0.149 | | 0.913 ± 0.109 | 0.991 ± 0.111 | 1.648 ± 0.148 | 1.177 ± 0.139 | 1.125 ± 0.150 |
| **9** | **12-;14-MeC28** | 0.819 ± 0.058 | | 0.933 ± 0.046 | 0.990 ± 0.049 | 1.203 ± 0.055 | 1.087 ± 0.054 | 0.972 ± 0.071 |
| **10** | **6-MeC28** | 0.819 ± 0.058 | | 0.933 ± 0.046 | 0.990 ± 0.049 | 1.203 ± 0.055 | 1.087 ± 0.054 | 0.972 ± 0.071 |
| **11** | **5-MeC28** | 0.409 ± 0.026 | | 0.461 ± 0.017 | 0.452 ± 0.027 | 0.537 ± 0.039 | 0.429 ± 0.023 | 0.407 ± 0.034 |
| **12** | **4-MeC28** | 0.208 ± 0.011 | | 0.227 ± 0.009 | 0.214 ± 0.009 | 0.223 ± 0.019 | 0.175 ± 0.011 | 0.164 ± 0.011 |
| **13** | **3-MeC28** | 0.721 ± 0.050 | | 0.852 ± 0.027 | 0.890 ± 0.036 | 1.047 ± 0.070 | 0.920 ± 0.031 | 0.879 ± 0.066 |
| **14** | **Unknow n** | 0.637 ± 0.048 | | 0.766 ± 0.022 | 0.799 ± 0.026 | 0.955 ± 0.064 | 0.888 ± 0.029 | 0.846 ± 0.059 |
| **15** | **unknown** | 0.259 ± 0.035 | | 0.269 ± 0.023 | 0.376 ± 0.038 | 0.453 ± 0.035 | 0.356 ± 0.035 | 0.320 ± 0.047 |
| **16** | **n-C29** | 9.903 ± 0.573 | | 10.882 ± 0.411 | 10.583 ± 0.528 | 10.227 ± 0.772 | 9.509 ± 0.400 | 10.676 ± 0.551 |
| **17** | **9-; 11-; 13-; 15-MeC29** | 15.862 ± 0.755 | | 17.972 ± 0.782 | 18.105 ± 0.924 | 20.442 ± 1.369 | 19.379 ± 1.017 | 18.194 ± 1.049 |
| **18** | **7-MeC29** | 3.411 ± 0.176 | | 3.810 ± 0.155 | 3.656 ± 0.152 | 4.512 ± 0.333 | 3.798 ± 0.210 | 3.821 ± 0.262 |
| **19** | **5-MeC29** | 8.496 ± 0.323 | | 9.676 ± 0.283 | 9.037 ± 0.335 | 9.229 ± 0.699 | 7.670 ± 0.414 | 7.488 ± 0.486 |
| **20** | **11,15-; 13,17-DimeC29** | 5.295 ± 0.360 | | 6.500 ± 0.259 | 7.191 ± 0.421 | 9.575 ± 0.701 | 8.448 ± 0.247 | 8.321 ± 0.487 |
| **21** | **7,11-DimeC29** | 1.166 ± 0.089 | | 1.309 ± 0.041 | 1.329 ± 0.048 | 1.831 ± 0.147 | 1.581 ± 0.089 | 1.529 ± 0.102 |
| **22** | **3-MeC29** | 9.643 ± 0.643 | | 11.457 ± 0.475 | 11.993 ± 0.636 | 11.805 ± 0.951 | 11.651 ± 0.749 | 12.068 ± 0.831 |
| **23** | **5,9-; 5,11-DimeC29** | 7.431 ± 0.486 | | 8.570 ± 0.207 | 8.675 ± 0.285 | 10.751 ± 0.891 | 9.163 ± 0.474 | 9.130 ± 0.648 |
| **24** | **3,7-; 3,9-;3,11-DimeC29** | 28.963 ± 2.039 | | 33.225 ± 0.960 | 32.973 ± 1.399 | 39.265 ± 2.707 | 35.263 ± 1.204 | 32.272 ± 1.874 |
| **25** | **7,11-DimeC30** | 3.930 ± 0.310 | | 4.610 ± 0.164 | 4.628 ± 0.209 | 5.106 ± 0.320 | 5.406 ± 0.142 | 4.574 ± 0.239 |
| **26** | **4,8-; 4,10-DimeC30** | 1.031 ± 0.088 | | 1.266 ± 0.033 | 1.187 ± 0.084 | 1.417 ± 0.170 | 1.467 ± 0.078 | 1.272 ± 0.089 |
| **27** | **Unknown** | 0.197 ± 0.020 | | 0.257 ± 0.015 | 0.243 ± 0.021 | 0.255 ± 0.019 | 0.293 ± 0.019 | 0.277 ± 0.010 |
| **28** | **11-; 13-; 15-MeC31** | 1.521 ± 0.108 | | 1.846 ± 0.129 | 1.789 ± 0.153 | 1.717 ± 0.132 | 1.743 ± 0.137 | 1.620 ± 0.082 |
| **29** | **11,15-; 13,17-DimeC31** | 0.314 ± 0.031 | | 0.418 ± 0.031 | 0.465 ± 0.031 | 0.615 ± 0.067 | 0.586 ± 0.018 | 0.609 ± 0.019 |
| **30** | **5,9-; 5,11-DimeC31** | 1.056 ± 0.076 | | 1.271 ± 0.037 | 1.125 ± 0.109 | 1.343 ± 0.189 | 1.368 ± 0.087 | 1.287 ± 0.096 |
| **31** | **10,12-DimeC32** | 0.351 ± 0.032 | | 0.434 ± 0.039 | 0.425 ± 0.048 | 0.400 ± 0.032 | 0.435 ± 0.040 | 0.355 ± 0.016 |
| **Peak No.** | **Compound** | **AD1-male** | | **AD2-male** | **AD3-male** | **AD4-male** | **AD5-male** | **AD6-male** |
| **1** | **n-C27** | 1.021 ± 0.083 | | 0.859 ± 0.074 | 1.081 ± 0.116 | 1.374 ± 0.105 | 2.202 ± 0.231 | 2.494 ± 0.209 |
| **2** | **11-; 13-MeC27** | 1.091 ± 0.110 | | 1.282 ± 0.125 | 2.100 ± 0.242 | 3.090 ± 0.195 | 4.851 ± 0.234 | 4.530 ± 0.257 |
| **3** | **5-MeC27** | 1.129 ± 0.104 | | 1.066 ± 0.112 | 1.511 ± 0.147 | 1.981 ± 0.133 | 2.859 ± 0.176 | 2.828 ± 0.179 |
| **4** | **11,15-DimeC27** | 0.158 ± 0.014 | | 0.188 ± 0.027 | 0.253 ± 0.026 | 0.354 ± 0.028 | 0.447 ± 0.023 | 0.474 ± 0.035 |
| **5** | **3-MeC27** | 1.435 ± 0.126 | | 1.404 ± 0.130 | 1.900 ± 0.190 | 2.533 ± 0.183 | 3.493 ± 0.215 | 3.605 ± 0.251 |
| **6** | **5, 9-; 5,11-DimeC27** | 0.625 ± 0.148 | | 0.735 ± 0.100 | 0.784 ± 0.077 | 1.097 ± 0.067 | 1.308 ± 0.163 | 1.054 ± 0.089 |
| **7** | **n-C28** | 0.650 ± 0.036 | | 0.576 ± 0.018 | 0.584 ± 0.022 | 0.661 ± 0.034 | 0.940 ± 0.060 | 0.956 ± 0.089 |
| **8** | **3, 9-; 3,11-DimeC27** | 0.638 ± 0.090 | | 0.578 ± 0.070 | 0.682 ± 0.057 | 0.843 ± 0.056 | 1.080 ± 0.058 | 1.018 ± 0.059 |
| **9** | **12-;14-MeC28** | 0.586 ± 0.037 | | 0.610 ± 0.022 | 0.713 ± 0.038 | 0.960 ± 0.039 | 1.385 ± 0.062 | 1.322 ± 0.067 |
| **10** | **6-MeC28** | 0.586 ± 0.037 | | 0.610 ± 0.022 | 0.713 ± 0.038 | 0.960 ± 0.039 | 1.297 ± 0.122 | 1.104 ± 0.128 |
| **11** | **5-MeC28** | 0.278 ± 0.009 | | 0.274 ± 0.009 | 0.279 ± 0.015 | 0.339 ± 0.020 | 0.467 ± 0.028 | 0.386 ± 0.044 |
| **12** | **4-MeC28** | 0.135 ± 0.005 | | 0.140 ± 0.003 | 0.148 ± 0.007 | 0.163 ± 0.007 | 0.271 ± 0.036 | 0.309 ± 0.069 |
| **13** | **3-MeC28** | 0.490 ± 0.016 | | 0.504 ± 0.017 | 0.515 ± 0.023 | 0.586 ± 0.033 | 0.699 ± 0.069 | 0.696 ± 0.042 |
| **14** | **Unknow n** | 0.434 ± 0.024 | | 0.432 ± 0.013 | 0.437 ± 0.018 | 0.507 ± 0.028 | 0.691 ± 0.023 | 0.545 ± 0.091 |
| **15** | **unknown** | 0.168 ± 0.029 | | 0.181 ± 0.017 | 0.197 ± 0.009 | 0.217 ± 0.012 | 0.278 ± 0.032 | 0.214 ± 0.014 |
| **16** | **n-C29** | 7.079 ± 0.356 | | 6.810 ± 0.261 | 6.505 ± 0.236 | 6.920 ± 0.404 | 8.835 ± 0.410 | 9.398 ± 1.068 |
| **17** | **9-; 11-; 13-; 15-MeC29** | 11.403 ± 0.326 | | 12.306 ± 0.306 | 14.069 ± 0.668 | 18.202 ± 0.868 | 23.902 ± 1.101 | 24.490 ± 1.829 |
| **18** | **7-MeC29** | 2.300 ± 0.078 | | 2.485 ± 0.057 | 2.738 ± 0.139 | 3.354 ± 0.153 | 4.490 ± 0.170 | 4.369 ± 0.316 |
| **19** | **5-MeC29** | 5.870 ± 0.210 | | 6.196 ± 0.159 | 6.161 ± 0.247 | 6.496 ± 0.300 | 8.082 ± 0.213 | 7.432 ± 0.707 |
| **20** | **11,15-; 13,17-DimeC29** | 3.849 ± 0.093 | | 4.437 ± 0.200 | 4.582 ± 0.236 | 5.457 ± 0.276 | 6.134 ± 0.212 | 6.233 ± 0.415 |
| **21** | **7,11-DimeC29** | 0.720 ± 0.027 | | 0.757 ± 0.033 | 0.739 ± 0.046 | 0.866 ± 0.051 | 1.116 ± 0.050 | 1.052 ± 0.055 |
| **22** | **3-MeC29** | 6.380 ± 0.372 | | 6.999 ± 0.256 | 7.081 ± 0.288 | 7.752 ± 0.508 | 9.456 ± 0.469 | 9.535 ± 0.967 |
| **23** | **5,9-; 5,11-DimeC29** | 5.106 ± 0.157 | | 5.433 ± 0.179 | 5.442 ± 0.263 | 5.863 ± 0.251 | 7.361 ± 0.255 | 6.675 ± 0.297 |
| **24** | **3,7-; 3,9-;3,11-DimeC29** | 20.088 ± 0.637 | | 20.249 ± 0.443 | 19.135 ± 0.872 | 19.747 ± 1.189 | 22.605 ± 1.007 | 19.629 ± 0.890 |
| **25** | **7,11-DimeC30** | 2.728 ± 0.078 | | 2.880 ± 0.111 | 2.750 ± 0.152 | 2.729 ± 0.171 | 2.997 ± 0.142 | 2.611 ± 0.092 |
| **26** | **4,8-; 4,10-DimeC30** | 0.694 ± 0.027 | | 0.676 ± 0.034 | 0.615 ± 0.046 | 0.593 ± 0.055 | 0.715 ± 0.046 | 0.506 ± 0.065 |
| **27** | **Unknown** | 0.133 ± 0.003 | | 0.140 ± 0.006 | 0.140 ± 0.007 | 0.151 ± 0.011 | 0.199 ± 0.009 | 0.161 ± 0.010 |
| **28** | **11-; 13-; 15-MeC31** | 0.965 ± 0.043 | | 1.313 ± 0.052 | 1.542 ± 0.117 | 1.875 ± 0.150 | 2.071 ± 0.104 | 2.019 ± 0.177 |
| **29** | **11,15-; 13,17-DimeC31** | 0.191 ± 0.010 | | 0.272 ± 0.014 | 0.283 ± 0.017 | 0.361 ± 0.028 | 0.402 ± 0.023 | 0.407 ± 0.039 |
| **30** | **5,9-; 5,11-DimeC31** | 0.625 ± 0.034 | | 0.670 ± 0.031 | 0.602 ± 0.052 | 0.614 ± 0.055 | 0.753 ± 0.041 | 0.587 ± 0.050 |
| **31** | **10,12-DimeC32** | 0.214 ± 0.012 | | 0.258 ± 0.012 | 0.277 ± 0.027 | 0.254 ± 0.025 | 0.282 ± 0.021 | 0.221 ± 0.020 |

The amount of different CHCs are shown as means ± SEM (ug/cockroach), GC peak numbers corresponded to Figure 1A and Pei et al., 2019 (Peak 22 and 23 were not separated in Pei et al., 2019). The writing of different compounds is in shorthand (Me: Methyl; Dime: Dimethyl; n: Normal; Unknown means the compound was not determined).
