## Supplementary material for "Modulation of fatty acid elongation sustains sexually dimorphic hydrocarbons and female attractiveness in *Blattella germanica* (L.)": S2Table

| **Quantification of individual cuticular hydrocarbons after RNAi of *BgElos*** | | | | | | | | | | | | | |
| --- | --- | --- | --- | --- | --- | --- | --- | --- | --- | --- | --- | --- | --- |
|  | **ds*Muslta*** | **ds*BgElo1*** | **ds*BgElo2*** | **ds*BgElo3*** | **ds*BgElo6*** | **ds*BgElo7*** | **ds*BgElo9*** | **ds*BgElo 10*** | **ds*BgElo 11*** | **ds*BgElo 14*** | **ds*BgElo 17*** | **ds*BgElo 20*** | **ds*BgElo 22*** |
| **n-C27** | 0.012 ± 0.005 | 0.016 ± 0.006 | 0.017 ± 0.006 | 0.014 ± 0.006 | 0.020 ± 0.009 | 0.029 ± 0.011 | 0.016 ± 0.005 | 0.013 ± 0.004 | 0.009 ± 0.002 | 0.013 ± 0.003 | 0.006 ± 0.001 | 0.018 ± 0.006 | 0.013 ± 0.005 |
| **11-; 13-MeC27** | 0.012 ± 0.006 | 0.027 ± 0.010 | 0.022 ± 0.010 | 0.016 ± 0.008 | 0.019 ± 0.007 | 0.030 ± 0.009 | 0.015 ± 0.005 | 0.023 ± 0.008 | 0.015 ± 0.005 | 0.026 ± 0.004 | 0.010 ± 0.003 | 0.016 ± 0.007 | 0.014 ± 0.006 |
| **5-MeC27** | 0.014 ± 0.005 | 0.018 ± 0.006 | 0.024 ± 0.008 | 0.016 ± 0.006 | 0.022 ± 0.008 | 0.030 ± 0.012 | 0.017 ± 0.005 | 0.015 ± 0.004 | 0.012 ± 0.003 | 0.016 ± 0.003 | 0.009 ± 0.002 | 0.019 ± 0.007 | 0.017 ± 0.006 |
| **11,15-DimeC27** | 0.002 ± 0.001 | 0.004 ± 0.001 | 0.004 ± 0.003 | 0.002 ± 0.001 | 0.003 ± 0.001 | 0.003 ± 0.001 | 0.002 ± 0.001 | 0.003 ± 0.001 | 0.002 ± 0.001 | 0.003 ± 0.001 | 0.001 ± 0.0005 | 0.002 ± 0.001 | 0.002 ± 0.001 |
| **3-MeC27** | 0.020 ± 0.006 | 0.024 ± 0.007 | 0.032 ± 0.009 | 0.026 ± 0.009 | 0.029 ± 0.010 | 0.043 ± 0.017 | 0.023 ± 0.008 | 0.021 ± 0.007 | 0.016 ± 0.004 | 0.021 ± 0.004 | 0.012 ± 0.003 | 0.026 ± 0.008 | 0.022 ± 0.008 |
| **5, 9-; 5,11-DimeC27** | 0.007 ± 0.006 | 0.036 ± 0.012 | 0.011 ± 0.006 | 0.009 ± 0.008 | 0.014 ± 0.006 | 0.014 ± 0.006 | 0.011 ± 0.006 | 0.032 ± 0.010 | 0.021 ± 0.006 | 0.030 ± 0.006 | 0.016 ± 0.005 | 0.011 ± 0.006 | 0.013 ± 0.007 |
| **n-C28** | 0.014 ± 0.003 | 0.026 ± 0.009 | 0.010 ± 0.002 | 0.009 ± 0.002 | 0.014 ± 0.005 | 0.012 ± 0.003 | 0.011 ± 0.002 | 0.026 ± 0.011 | 0.016 ± 0.004 | 0.020 ± 0.004 | 0.011 ± 0.003 | 0.011 ± 0.001 | 0.009 ± 0.001 |
| **3, 9-; 3,11-DimeC27** | 0.004 ± 0.001 | 0.019 ± 0.005 | 0.017 ± 0.009 | 0.011 ± 0.005 | 0.012 ± 0.005 | 0.020 ± 0.006 | 0.011 ± 0.005 | 0.019 ± 0.007 | 0.012 ± 0.002 | 0.015 ± 0.003 | 0.008 ± 0.002 | 0.014 ± 0.005 | 0.012 ± 0.007 |
| **12-;14-MeC28** | 0.010 ± 0.001 | 0.016 ± 0.004 | 0.013 ± 0.002 | 0.012 ± 0.002 | 0.013 ± 0.002 | 0.014 ± 0.002 | 0.011 ± 0.002 | 0.014 ± 0.004 | 0.011 ± 0.002 | 0.012 ± 0.002 | 0.007 ± 0.001 | 0.012 ± 0.002 | 0.010 ± 0.002 |
| **6-MeC28** | 0.004 ± 0.001 | 0.005 ± 0.001 | 0.006 ± 0.001 | 0.006 ± 0.001 | 0.006 ± 0.001 | 0.006 ± 0.002 | 0.005 ± 0.001 | 0.004 ± 0.001 | 0.004 ± 0.0004 | 0.003 ± 0.0004 | 0.002 ± 0.001 | 0.006 ± 0.001 | 0.005 ± 0.001 |
| **5-MeC28** | 0.002 ± 0.0003 | 0.002 ± 0.0005 | 0.003 ± 0.0004 | 0.002 ± 0.0005 | 0.002 ± 0.001 | 0.003 ± 0.001 | 0.003 ± 0.0003 | 0.001 ± 0.0003 | 0.002 ± 0.0002 | 0.001 ± 0.0002 | 0.001 ± 0.0002 | 0.003 ± 0.0003 | 0.002 ± 0.0003 |
| **4-MeC28** | 0.008 ± 0.001 | 0.010 ± 0.002 | 0.012 ± 0.001 | 0.010 ± 0.002 | 0.010 ± 0.002 | 0.010 ± 0.002 | 0.009 ± 0.001 | 0.008 ± 0.002 | 0.007 ± 0.001 | 0.006 ± 0.001 | 0.005 ± 0.001 | 0.010 ± 0.001 | 0.009 ± 0.001 |
| **3-MeC28** | 0.003 ± 0.001 | 0.007 ± 0.001 | 0.011 ± 0.001 | 0.010 ± 0.001 | 0.007 ± 0.003 | 0.009 ± 0.002 | 0.009 ± 0.001 | 0.005 ± 0.001 | 0.005 ± 0.000 | 0.005 ± 0.001 | 0.003 ± 0.001 | 0.009 ± 0.001 | 0.008 ± 0.001 |
| **Unknow n** | 0.006 ± 0.001 | 0.003 ± 0.001 | 0.002 ± 0.0003 | 0.002 ± 0.001 | 0.004 ± 0.002 | 0.002 ± 0.0003 | 0.002 ± 0.0003 | 0.003 ± 0.001 | 0.002 ± 0.0005 | 0.002 ± 0.0004 | 0.002 ± 0.0004 | 0.002 ± 0.0003 | 0.002 ± 0.0003 |
| **unknown** | 0.004 ± 0.002 | 0.005 ± 0.002 | 0.005 ± 0.001 | 0.004 ± 0.001 | 0.005 ± 0.002 | 0.005 ± 0.002 | 0.004 ± 0.001 | 0.005 ± 0.002 | 0.003 ± 0.001 | 0.004 ± 0.001 | 0.002 ± 0.001 | 0.005 ± 0.002 | 0.004 ± 0.002 |
| **n-C29** | 0.125 ± 0.014 | 0.132 ± 0.031 | 0.101 ± 0.015 | 0.102 ± 0.019 | 0.123 ± 0.010 | 0.117 ± 0.030 | 0.111 ± 0.019 | 0.094 ± 0.015 | 0.106 ± 0.017 | 0.106 ± 0.023 | 0.101 ± 0.022 | 0.116 ± 0.012 | 0.108 ± 0.011 |
| **9-; 11-; 13-; 15-MeC29** | 0.189 ± 0.021 | 0.250 ± 0.048 | 0.204 ± 0.027 | 0.214 ± 0.031 | 0.204 ± 0.012 | 0.182 ± 0.026 | 0.180 ± 0.024 | 0.199 ± 0.030 | 0.222 ± 0.025 | 0.214 ± 0.040 | 0.200 ± 0.026 | 0.174 ± 0.019 | 0.188 ± 0.017 |
| **7-MeC29** | 0.039 ± 0.005 | 0.054 ± 0.011 | 0.046 ± 0.006 | 0.048 ± 0.008 | 0.043 ± 0.003 | 0.043 ± 0.008 | 0.038 ± 0.005 | 0.042 ± 0.007 | 0.046 ± 0.006 | 0.044 ± 0.007 | 0.039 ± 0.006 | 0.040 ± 0.004 | 0.042 ± 0.004 |
| **5-MeC29** | 0.099 ± 0.011 | 0.101 ± 0.026 | 0.102 ± 0.016 | 0.087 ± 0.014 | 0.101 ± 0.007 | 0.089 ± 0.020 | 0.086 ± 0.011 | 0.086 ± 0.014 | 0.095 ± 0.013 | 0.083 ± 0.015 | 0.087 ± 0.013 | 0.092 ± 0.012 | 0.097 ± 0.013 |
| **11,15-; 13,17-DimeC29** | 0.061 ± 0.012 | 0.100 ± 0.022 | 0.077 ± 0.009 | 0.068 ± 0.011 | 0.072 ± 0.007 | 0.055 ± 0.008 | 0.063 ± 0.008 | 0.073 ± 0.020 | 0.073 ± 0.014 | 0.069 ± 0.011 | 0.072 ± 0.009 | 0.054 ± 0.008 | 0.070 ± 0.011 |
| **7,11-DimeC29** | 0.015 ± 0.001 | 0.015 ± 0.003 | 0.017 ± 0.002 | 0.017 ± 0.003 | 0.016 ± 0.002 | 0.012 ± 0.002 | 0.014 ± 0.002 | 0.014 ± 0.003 | 0.015 ± 0.002 | 0.013 ± 0.002 | 0.012 ± 0.002 | 0.014 ± 0.003 | 0.015 ± 0.002 |
| **3-MeC29** | 0.126 ± 0.008 | 0.101 ± 0.019 | 0.097 ± 0.013 | 0.096 ± 0.016 | 0.095 ± 0.006 | 0.078 ± 0.016 | 0.079 ± 0.008 | 0.084 ± 0.010 | 0.097 ± 0.013 | 0.083 ± 0.014 | 0.090 ± 0.010 | 0.083 ± 0.010 | 0.112 ± 0.018 |
| **5,9-; 5,11-DimeC29** | 0.093 ± 0.006 | 0.136 ± 0.026 | 0.131 ± 0.018 | 0.130 ± 0.022 | 0.128 ± 0.007 | 0.105 ± 0.022 | 0.107 ± 0.011 | 0.113 ± 0.014 | 0.130 ± 0.018 | 0.112 ± 0.019 | 0.122 ± 0.014 | 0.112 ± 0.014 | 0.095 ± 0.011 |
| **3,7-; 3,9-;3,11-DimeC29** | 0.361 ± 0.027 | 0.348 ± 0.066 | 0.411 ± 0.054 | 0.391 ± 0.059 | 0.389 ± 0.044 | 0.297 ± 0.055 | 0.308 ± 0.029 | 0.351 ± 0.049 | 0.374 ± 0.047 | 0.325 ± 0.051 | 0.359 ± 0.038 | 0.339 ± 0.036 | 0.336 ± 0.030 |
| **7,11-DimeC30** | 0.050 ± 0.006 | 0.044 ± 0.014 | 0.063 ± 0.011 | 0.056 ± 0.012 | 0.057 ± 0.011 | 0.040 ± 0.010 | 0.046 ± 0.004 | 0.046 ± 0.009 | 0.052 ± 0.009 | 0.040 ± 0.007 | 0.044 ± 0.007 | 0.049 ± 0.008 | 0.049 ± 0.008 |
| **4,8-; 4,10-DimeC30** | 0.015 ± 0.001 | 0.009 ± 0.003 | 0.019 ± 0.004 | 0.016 ± 0.004 | 0.015 ± 0.003 | 0.009 ± 0.003 | 0.013 ± 0.001 | 0.009 ± 0.003 | 0.011 ± 0.003 | 0.008 ± 0.002 | 0.010 ± 0.002 | 0.014 ± 0.003 | 0.012 ± 0.002 |
| **Unknown** | 0.004 ± 0.001 | 0.002 ± 0.001 | 0.004 ± 0.001 | 0.003 ± 0.001 | 0.003 ± 0.0004 | 0.002 ± 0.001 | 0.003 ± 0.001 | 0.002 ± 0.0004 | 0.002 ± 0.0003 | 0.002 ± 0.0003 | 0.001 ± 0.0003 | 0.003 ± 0.0004 | 0.002 ± 0.0003 |
| **11-; 13-; 15-MeC31** | 0.019 ± 0.002 | 0.016 ± 0.006 | 0.019 ± 0.003 | 0.020 ± 0.006 | 0.016 ± 0.002 | 0.011 ± 0.003 | 0.016 ± 0.003 | 0.011 ± 0.002 | 0.020 ± 0.004 | 0.013 ± 0.003 | 0.017 ± 0.003 | 0.015 ± 0.002 | 0.016 ± 0.003 |
| **11,15-; 13,17-DimeC31** | 0.005 ± 0.001 | 0.006 ± 0.002 | 0.005 ± 0.001 | 0.005 ± 0.001 | 0.004 ± 0.001 | 0.003 ± 0.001 | 0.004 ± 0.001 | 0.003 ± 0.001 | 0.004 ± 0.001 | 0.003 ± 0.001 | 0.003 ± 0.001 | 0.004 ± 0.001 | 0.004 ± 0.001 |
| **5,9-; 5,11-DimeC31** | 0.013 ± 0.002 | 0.007 ± 0.003 | 0.016 ± 0.002 | 0.015± 0.004 | 0.013 ± 0.002 | 0.008 ± 0.002 | 0.012 ± 0.002 | 0.007 ± 0.002 | 0.010 ± 0.003 | 0.006 ± 0.001 | 0.009 ± 0.002 | 0.012 ± 0.002 | 0.011 ± 0.002 |
| **10,12-DimeC32** | 0.005 ± 0.001 | 0.002 ± 0.001 | 0.006 ± 0.002 | 0.005 ± 0.001 | 0.005 ± 0.001 | 0.003 ± 0.001 | 0.004 ± 0.001 | 0.002 ± 0.001 | 0.003 ± 0.001 | 0.002 ± 0.000 | 0.003 ± 0.001 | 0.005 ± 0.001 | 0.004 ± 0.001 |

The amount of different CHCs are shown as means ± SEM (ug/cockroach). The writing of different compounds is in shorthand (Me: Methyl; Dime: Dimethyl; n: Normal; Unknown means the compound was not determined).
