## Supplementary material for "Modulation of fatty acid elongation sustains sexually dimorphic hydrocarbons and female attractiveness in *Blattella germanica* (L.)": S3Table

**Calculation of individual fatty acid methyl ester proportion in yeast expression.**

|  | **GFP** | **GFP**  **+C20** | **GFP**  **+C22** | **GFP +C24** | **GFP +C26** | **GFP +C28** | **GFP +2-MeC16** | **GFP +14-MeC16** |
| --- | --- | --- | --- | --- | --- | --- | --- | --- |
| **C10:0** | 0.017 ±  0.012 | 0.130 ±  0.111 | 0.013 ±  0.005 | 0.010 ±  0.001 | 0.013 ±  0.006 | 0.013 ±  0.006 | 0.020 ±  0.010 | 0.025 ±  0.015 |
| **C12:0** | 0.100 ±  0.020 | 0.143 ±  0.082 | 0.125 ±  0.061 | 0.047 ±  0.006 | 0.093 ±  0.084 | 0.083 ±  0.009 | 0.080 ±  0.035 | 0.095 ±  0.045 |
| **C14:1** | 0.033 ±  0.006 | 0.173 ±  0.125 | 0.033 ±  0.013 | 0.020 ±  0.002 | 0.027 ±  0.012 | 0.023 ±  0.005 | 0.037 ±  0.023 | 0.040 ±  0.020 |
| **C14:0** | 0.263 ±  0.023 | 0.403 ±  0.096 | 0.310 ±  0.146 | 0.153 ±  0.021 | 0.257 ±  0.212 | 0.230 ±  0.025 | 0.223 ±  0.098 | 0.265 ±  0.095 |
| **C15:1** | 0.027 ±  0.015 | 0.070 ±  0.044 | 0.020 ±  0.008 | 0.010 ±  0.001 | 0.017 ±  0.012 | 0.013 ±  0.005 | 0.023 ±  0.012 | 0.015 ±  0.005 |
| **C15:0** | 0.043 ±  0.015 | 0.047 ±  0.017 | 0.040 ±  0.016 | 0.017 ±  0.006 | 0.030 ±  0.026 | 0.010 ±  0.001 | 0.037 ±  0.023 | 0.035 ±  0.015 |
| **C16:1** | 54.173 ±  6.146 | 54.293 ±  3.193 | 50.735 ±  2.608 | 55.927 ±  4.134 | 52.343 ±  8.336 | 53.823 ±  3.908 | 52.213 ±  1.807 | 56.645 ±  2.205 |
| **C16:0** | 22.500 ±  3.146 | 21.340 ±  2.405 | 22.048 ±  2.473 | 20.670 ±  3.072 | 21.490 ±  3.132 | 21.640 ±  1.447 | 21.823 ±  0.098 | 20.595 ±  0.345 |
| **C17:1** | 0.037 ±  0.013 | 0.053 ±  0.009 | 0.048 ±  0.015 | 0.063 ±  0.006 | 0.043 ±  0.006 | 0.050 ±  0.011 | 0.023 ±  0.012 | 0.065 ±  0.005 |
| **C17:0** | 0.023 ±  0.012 | 0.067 ±  0.010 | 0.033 ±  0.005 | 0.043 ±  0.006 | 0.033 ±  0.006 | 0.053 ±  0.079 | 0.017 ±  0.006 | 0.015 ±  0.005 |
| **C18:1** | 20.020 ±  4.812 | 18.940 ±  1.283 | 20.718 ±  2.250 | 18.790 ±  0.506 | 19.153 ±  1.880 | 18.808 ±  0.829 | 20.480 ±  0.485 | 18.210 ±  0.710 |
| **C18:0** | 2.263 ±  0.473 | 2.380 ±  0.354 | 3.583 ±  0.735 | 3.577 ±  0.802 | 3.377 ±  0.664 | 3.377 ±  0.192 | 4.057 ±  0.318 | 3.400 ±  0.360 |
| **C20:1** | Trace | 0.010 ±  0.001 | 0.010 ±  0.008 | Trace | Trace | Trace | Trace’ | Trace |
| **C20:0** | 0.030 ±  0.017 | 1.420 ±  0.888 | 0.063 ±  0.060 | 0.010 ±  0.001 | 0.017 ±  0.006 | 0.027 ±  0.001 | 0.023 ±  0.012 | 0.015 ±  0.005 |
| **C22:1** | Trace | 0.017 ±  0.006 | 0.010 ±  0.014 | Trace | Trace | Trace | 0.007 ±  0.006 | Trace |
| **C22:0** | 0.010 ±  0.001 | 0.017 ±  0.006 | 1.548 ±  0.425 | 0.010 ±  0.006 | 0.012 ±  0.006 | 0.013 ±  0.007 | 0.013 ±  0.012 | 0.010 ±  0.006 |
| **C24:0** | 0.023 ±  0.005 | 0.047 ±  0.012 | 0.040 ±  0.011 | 0.047 ±  0.012 | 0.031 ±  0.008 | 0.020 ±  0.007 | 0.020 ±  0.007 | 0.015 ±  0.005 |
| **C26:0** | 0.427 ±  0.105 | 0.433 ±  0.051 | 0.613 ±  0.051 | 0.647 ±  0.073 | 3.073 ±  0.535 | 0.297 ±  0.042 | 0.863 ±  0.068 | 0.545 ±  0.075 |
| **C28:0** | 0.012 ±  0.001 | 0.010 ±  0.001 | 0.023 ±  0.005 | 0.007 ±  0.006 | 0.017 ±  0.006 | 1.523 ±  0.589 | 0.023 ±  0.012 | 0.015 ±  0.005 |
| **C30:0** | NF | NF | NF | NF | NF | NF | NF | NF |

|  | **BgElo12** | **BgElo12**  **+C20** | **BgElo12**  **+C22** | **BgElo12**  **+C24** | **BgElo12**  **+C26** | **BgElo12**  **+C28** | **BgElo12+2-MeC16** | **BgElo12+14-MeC16** |
| --- | --- | --- | --- | --- | --- | --- | --- | --- |
| **C10:0** | 0.022 ±  0.010 | 0.033 ±  0.015 | 0.023 ±  0.005 | 0.020 ±  0.01 | 0.023 ±  0.006 | 0.060 ±  0.005 | 0.023 ±  0.0153 | 0.013 ±  0.006 |
| **C12:0** | 0.084 ± 0.025 | 0.110 ±  0.030 | 0.103 ±  0.025 | 0.107 ±  0.035 | 0.110 ±  0.030 | 0.230 ±  0.088 | 0.110 ±  0.050 | 0.083 ±  0.015 |
| **C14:1** | 0.038 ± 0.012 | 0.053 ±  0.015 | 0.053 ±  0.015 | 0.053 ±  0.015 | 0.053 ±  0.015 | 0.148 ±  0.012 | 0.060 ±  0.030 | 0.040 ±  0.010 |
| **C14:0** | 0.370 ±  0.054 | 0.453 ±  0.095 | 0.420 ±  0.090 | 0.433 ±  0.110 | 0.430 ±  0.101 | 0.428 ±  0.157 | 0.343 ±  0.135 | 0.277 ±  0.015 |
| **C15:1** | 0.014 ±  0.004 | 0.023 ±  0.006 | 0.020 ±  0.010 | 0.020 ±  0.010 | 0.017 ±  0.005 | 0.035 ±  0.005 | 0.037 ±  0.015 | 0.023 ±  0.005 |
| **C15:0** | 0.0275 ±  0.008 | 0.041 ±  0.010 | 0.040 ±  0.010 | 0.030 ±  0.010 | 0.038 ±  0.010 | 0.058 ±  0.010 | 0.057 ±  0.025 | 0.033 ±  0.015 |
| **C16:1** | 52.600 ±  2.482 | 49.413 ±  0.255 | 48.413 ±  0.425 | 49.533 ±1.255 | 47.699 ±  0.595 | 52.263 ±  4.181 | 50.027 ±  3.015 | 57.580 ±  1.040 |
| **C16:0** | 22.087 ±  2.913 | 22.823 ±  1.393 | 22.760 ±  1.650 | 23.021 ±1.905 | 22.957 ±  2.105 | 21.689 ±  2.688 | 23.697 ±  1.285 | 19.557 ±  0.320 |
| **C17:1** | 0.018 ±  0.004 | 0.025 ±  0.005 | 0.023 ±  0.006 | 0.029 ±  0.003 | 0.033 ±  0.006 | 0.020 ±  0.001 | 0.030 ±  0.010 | 0.053 ±  0.043 |
| **C17:0** | 0.010 ±  0.002 | 0.010 ±  0.002 | 0.010 ±  0.002 | 0.010 ±  0.001 | 0.010 ±  0.001 | 0.041 ±  0.0020 | 0.020 ±  0.010 | 0.013 ±  0.004 |
| **C18:1** | 20.500 ±  0.669 | 20.760 ±  0.860 | 20.483 ±  0.185 | 21.173 ±  0.375 | 20.907 ±  0.375 | 19.083 ±  1.2336 | 20.240 ±  0.310 | 18.587 ±  1.346 |
| **C18:0** | 3.596 ±  0.509 | 4.363 ±  0.375 | 4.370 ±  0.190 | 4.323 ±  0.235 | 4.377 ±  0.295 | 4.463 ±  0.466 | 4.550 ±  0.630 | 3.010 ±  0.192 |
| **C20:1** | Trace | 0.010 ±  0.001 | 0.010 ±  0.002 | Trace | Trace | Trace | 0.010 ±  0.010 | Trace |
| **C20:0** | 0.017 ±  0.004 | 0.910 ±  0.470 | 0.030 ±  0.002 | 0.020 ±  0.001 | 0.020 ±  0.001 | 0.023 ±  0.003 | 0.020 ±  0.010 | 0.023 ±  0.005 |
| **C22:1** | Trace | Trace | Trace | Trace | Trace | 0.010 ±  0.001 | Trace | Trace |
| **C22:0** | 0.010 ±  0.002 | 0.010 ±  0.001 | 2.207 ±  0.915 | 0.010 ±  0.001 | 0.010 ±  0.001 | 0.035 ±  0.004 | 0.010 ±  0.010 | 0.010 ±  0.001 |
| **C24:0** | 0.011 ±  0.004 | 0.010 ±  0.001 | 0.030 ±  0.001 | 0.250 ±  0.120 | 0.017 ±  0.006 | Trace | 0.013 ±  0.006 | 0.027 ±  0.002 |
| **C26:0** | 0.661 ±  0.249 | 0.930 ±  0.120 | 0.980 ±  0.110 | 0.930 ±  0.110 | 3.277 ±  0.705 | 0.113 ±  0.016 | 0.707 ±  0.445 | 0.630 ±  0.131 |
| **C28:0** | 0.012 ±  0.006 | 0.020 ±  0.005 | 0.030 ±  0.010 | 0.027 ±  0.005 | 0.030 ±  0.001 | 1.208 ±  0.189 | 0.040 ±  0.020 | 0.030 ±  0.017 |
| **C30:0** | NF | NF | NF | NF | NF | 0.090 ±  0.005 | NF | NF |

|  | **BgElo24** | **BgElo24**  **+C20** | **BgElo24**  **+C22** | **BgElo24**  **+C24** | **BgElo24**  **+C26** | **BgElo24**  **+C28** | **BgElo24**  **+2-MeC16** | **BgElo24**  **+14-MeC16** |
| --- | --- | --- | --- | --- | --- | --- | --- | --- |
| **C10:0** | 0.026 ±  0.005 | 0.037 ±  0.012 | 0.030 ±  0.010 | 0.033 ±  0.012 | 0.038 ±  0.021 | 0.017 ±  0.012 | 0.018 ±  0.008 | 0.023 ±  0.005 |
| **C12:0** | 0.098 ±  0.025 | 0.140 ±  0.044 | 0.100 ±  0.030 | 0.140 ±  0.044 | 0.133 ±  0.055 | 0.087 ±  0.029 | 0.082 ±  0.018 | 0.093 ±  0.019 |
| **C14:1** | 0.052 ±  0.011 | 0.083 ±  0.022 | 0.060 ±  0.011 | 0.087 ±  0.013 | 0.073 ±  0.012 | 0.060 ±  0.007 | 0.046 ±  0.004 | 0.055 ±  0.010 |
| **C14:0** | 0.420 ±  0.064 | 0.583 ±  0.171 | 0.427 ±  0.085 | 0.530 ±  0.049 | 0.498 ±  0.075 | 0.360 ±  0.054 | 0.226 ±  0.047 | 0.260 ±  0.051 |
| **C15:1** | 0.022 ±  0.010 | 0.037 ±  0.012 | 0.020 ±  0.010 | 0.033 ±  0.012 | 0.030 ±  0.012 | 0.023 ±  0.006 | 0.030 ±  0.010 | 0.023 ±  0.005 |
| **C15:0** | 0.042 ±  0.010 | 0.053 ±  0.013 | 0.040 ±  0.010 | 0.050 ±  0.007 | 0.045 ±  0.007 | 0.033 ±  0.006 | 0.038 ±  0.008 | 0.028 ±  0.002 |
| **C16:1** | 55.68 ±  2.923 | 51.755 ±  2.475 | 55.567 ±  1.635 | 52.570 ±  2.154 | 52.040 ± 2.863 | 51.587 ±  2.567 | 55.792 ±  3.567 | 54.583 ±  1.866 |
| **C16:0** | 20.662 ±  1.737 | 21.237 ±  2.042 | 19.180 ±  1.140 | 22.183 ±  1.121 | 20.750 ±  2.124 | 20.803 ±  1.558 | 20.142 ±  1.086 | 20.928 ±  2.102 |
| **C17:1** | 0.026 ±  0.005 | 0.027 ±  0.002 | 0.017 ±  0.002 | 0.027 ±  0.003 | 0.025 ±  0.003 | 0.023 ±  0.004 | 0.026 ±  0.005 | 0.070 ±  0.018 |
| **C17:0** | 0.010 ±  0.001 | 0.013 ±  0.006 | 0.010 ±  0.001 | 0.017 ±  0.006 | 0.015 ±  0.006 | 0.010 ±  0.001 | 0.014 ±  0.005 | 0.013 ±  0.005 |
| **C18:1** | 18.046 ±  0.626 | 19.217 ±  0.815 | 18.067 ±  0.585 | 18.213 ±  0.596 | 18.745 ±  1.063 | 19.970 ±  0.227 | 18.022 ±  2.315 | 18.653 ±  1.313 |
| **C18:0** | 3.648 ±  0.487 | 4.050 ±  0.301 | 3.557 ±  0.175 | 4.267 ±  0.292 | 3.653 ±  0.684 | 4.277 ±  0.356 | 4.200 ±  0.600 | 3.755 ±  0.339 |
| **C20:1** | Trace | 0.013 ±  0.006 | 0.010 ±  0.001 | Trace | 0.011 ±  0.001 | Trace | 0.022 ±  0.016 | Trace |
| **C20:0** | 0.052 ±  0.019 | 0.552 ±  0.054 | 0.040 ±  0.001 | 0.037 ±  0.006 | 0.043 ±  0.032 | 0.040 ±  0.002 | 0.028 ±  0.008 | 0.030 ±  0.012 |
| **C22:1** | 0.016 ±  0.003 | 0.012 ±  0.001 | 0.010 ±  0.001 | Trace | Trace | 0.020 ±  0.001 | Trace | Trace |
| **C22:0** | 0.018 ±  0.008 | 0.023 ±  0.006 | 0.960 ±  0.550 | 0.083 ±  0.010 | 0.015 ±  0.006 | 0.017 ±  0.006 | 0.012 ±  0.004 | 0.015 ±  0.006 |
| **C24:0** | 0.016 ±  0.005 | 0.027 ±  0.006 | 0.020 ±  0.001 | 0.217 ±  0.039 | 0.018 ±  0.005 | 0.020 ±  0.003 | 0.014 ±  0.005 | 0.015 ±  0.006 |
| **C26:0** | 0.312 ±  0.081 | 0.513 ±  0.127 | 0.437 ±  0.035 | 0.387 ±  0.105 | 2.853 ±  0.382 | 0.443 ±  0.057 | 0.232 ±  0.110 | 0.278 ±  0.090 |
| **C28:0** | 0.570 ±  0.194 | 1.043 ±  0.490 | 0.913 ±  0.035 | 0.727 ±  0.280 | 0.665 ±  0.272 | 1.527 ±  0.808 | 0.638 ±  0.201 | 0.768 ±  0.140 |
| **C30:0** | 0.280 ±  0.141 | 0.583 ±  0.188 | 0.537 ±  0.115 | 0.387 ±  0.258 | 0.343 ±  0.276 | 0.690 ±  0.106 | 0.420 ±  0.157 | 0.400 ±  0.065 |

The proportions of different FAMEs are shown as means ± SD (%) and calculated from 3–5 independent replicates (single colony), different FAMEs are represented by their corresponding FAs. Trace represents the proportion lower than 0.01%. GFP+C20 represents the yeast carried with pYES2-GFP recombination vector, and the medium added with C20:0 fatty acid, and so on.
