## Supplementary material for "Modulation of fatty acid elongation sustains sexually dimorphic hydrocarbons and female attractiveness in *Blattella germanica* (L.)": S4Table

Gene specific primers used in this research.

| Products (bp) | Forward (5' to 3') | Reverse (5' to 3') | Purpose |
| --- | --- | --- | --- |
| *BgElo1* (940) | ATGACAGCTGTAATCAGAG | CTAAGAATTAGGTCCCTATTAATC | CDS cloning |
| *BgElo2* (1041) | ATCATGGCTCAACTCGTGCG | GCCCTGCTGAAATGAATCGT | CDS cloning |
| *BgElo3* (1115) | GCCGACACATAAACTCTCCGT | GGCACCCATTGAAAGCCATC | CDS cloning |
| *BgElo4* (560) | GCAGACCCTCGAGTGAACAA | AGGTGCCTCTTCCACCAAAG | CDS cloning |
| *BgElo5* (942) | ATGCATAACTGGAGAGTAAAAGAC | TTATCCAGTCTTTTTATCACCTTCC | CDS cloning |
| *BgElo6* (890) | ATGCTGCCACAGCTCAGTAA | TCAGGCATTTCTGCACGATCT | CDS cloning |
| *BgElo7* (833) | TGGTTCCTCATCACAAGTCCG | TGGCTCTCATCAGTGCAAGG | CDS cloning |
| *BgElo8* (806) | TCACAATGTCAAATCTTATGCATC | CAGGTGCTATCACACTGCCT | CDS cloning |
| *BgElo9* (1023) | GCTCAACAAAATGGCGCAGA | GTTCCAATCTGTCTACGACGATAC | CDS cloning |
| *BgElo10* (1062) | ACTGAGTGCCCCGAAGAACA | GTGATGACTGGGGTTTTCCCG | CDS cloning |
| *BgElo11* (849) | CCAGGGCTGCCATCCTAACT | TGTGGAAAGACTGAACTAAGCCA | CDS cloning |
| *BgElo12* (887) | GGGAACATCACAGCTCTCAGTA | TTATTGCTTATGAGATTTTC | CDS cloning |
| *BgElo13* (986) | AAACTTGACAACGACACCGC | TGGTCTAAAATATTGTGGCCCTATT | CDS cloning |
| *BgElo14* (979) | GGTTTCCGTCCTCACCTATTCA | TACCCAAACGGATCACCCTG | CDS cloning |
| *BgElo15* (900) | ATGGCAACCTTGATAAACGAAACT | TCCGACTGCATTTGCTACTCT | CDS cloning |
| *BgElo16* (388) | CACCATGGATCCCCGTCTTC | CTGATCCACAACCAACCAATCA | CDS cloning |
| *BgElo17* (881) | ATGGTCTCCTGGATCAATGCC | GCTCGATGTAATGTCAATATGGTT | CDS cloning |
| *BgElo18* (195*)* | CCAGGTGGTCATGGAATGCT | TTTCGAGACAAAAAGCGTCG | CDS cloning |
| *BgElo19* (331) | AAGTGCCCTGACTCCAGTTG | GGTTCGCAGATCAGGCTGTA | CDS cloning |
| *BgElo20* (994) | GATCTGATCCACCTGTCGGC | TGGTTTATCCCTGCTACTCATCT | CDS cloning |
| *BgElo21* (345) | ACTACATCAGCAATGCCACGA | GCCAGCATGTCTCTGCTATCT | CDS cloning |
| *BgElo22* (1345) | GGGTCAGAGGTTTCGCACTT | CACAAACTCATGGACCCCCA | CDS cloning |
| *BgElo23* (303) | TGTGTTCCGTGATCAGCCAT | AGTCCAAGAGTCGTACTTGCG | CDS cloning |
| *BgElo24* (958) | TTTCAGCGGGACTCGTCATC | TGACCGCTTTTGAACGCAAT | CDS cloning |
| *BgElo1* (104) | TCAGAGCTTGCTTCAAGGGA | CACGGTTACTGGTTGCCAAA | RT-qPCR, *E* = 99.2% |
| *BgElo2* (86) | CACAGCACATTCTTTGCCCT | TATCTTGGGCCCATTGCAGA | RT-qPCR, *E* = 100.8% |
| *BgElo3* (113) | ATTCACACCAGGTGGCCATA | AGAACTTCTGGAAGCGAGGT | RT-qPCR, *E* = 99.3% |
| *BgElo4* (139) | CCATCACTCAGTCACACCCA | AGGACCCATTGCTGATACCAT | RT-qPCR, *E* = 97.7% |
| *BgElo5* (93) | TGCCTGGAGGTCAAGCTTTT | CTGGACCCAGTGCAGAAAGA | RT-qPCR, *E* = 98.5% |
| *BgElo6* (113) | TGGCGTCGAGAAAACCCTAT | TGCAAGCATCCCTCCTGTAA | RT-qPCR, *E* = 97.7% |
| *BgElo7* (78) | GCACATGTATCACCACACCG | AAAGTACCATGGCCTCCTGG | RT-qPCR, *E* = 100.5% |
| *BgElo8* (129) | TCCTGATGAGCAGTCCATACC | TGTACGCCACAAGAACTCTCT | RT-qPCR, *E* = 96.3% |
| *BgElo9* (106) | CAACTATCCTCGGGCGTTTG | TGCTTTCCCCCTCTGCTTAT | RT-qPCR, *E* = 100.0% |
| *BgElo10* (110) | ACATGGCCTCTCATGGACAA | CTGTACGGTTTTCGCTGGTC | RT-qPCR, *E* = 101.1% |
| *BgElo11* (80) | TCAGTTCTGCCTAGCTGCTT | ATACGGCAACACCTCTTGGA | RT-qPCR, *E* = 101.5% |
| *BgElo12* (80) | AAATTACGCTTGGATGCTCTCC | TGTCCACCACAATAAAGATGCC | RT-qPCR, *E* = 96.4% |
| *BgElo13* (92) | GAGTTGCTCAGTGCTTCTACG | TTTGCTATCCTCAGCTCGTCT | RT-qPCR, *E* = 95.8% |
| *BgElo14* (107) | TGTCGGTAAATCAGATCCAAGG | GACCCCAGCTCAAAACGAAA | RT-qPCR, *E* = 98.2% |
| *BgElo15* (84) | GAGTGTCGCCAGATATGTTCC | TACCCGTACATGAAAACGTGC | RT-qPCR, *E* = 97.7% |
| *BgElo16* (117) | TTCTGCAAATCGCTGCATGT | GCTGCATTGTCTCCGGATTT | RT-qPCR, *E* = 102.7% |
| *BgElo17* (101) | ACGACGACATGGCCAGAATA | GGCAACGTGTGCAAAAACAC | RT-qPCR, *E* = 101.4% |
| *BgElo18* (109) | GGCCAAATCTCATCAGTGCC | TTCGAGACAAAAAGCGTCGT | RT-qPCR, *E* = 98.8% |
| *BgElo19* (99) | CATGAAGGACAAACCAGCGT | CGAAAGGCCCTGATCACAAT | RT-qPCR, *E* =98.3% |
| *BgElo20* (114) | ACATCACCACCCTCCAGATG | CGTTGATGCCCAGAAGGAAG | RT-qPCR, *E* = 103.5% |
| *BgElo21* (103) | CAGACCTGCCTACAACCTGA | CCTAAATGTCAGAACGGCGG | RT-qPCR, *E* = 97.8% |
| *BgElo22* (110) | TCATCGTCCTACGGAAGCAG | CCACCGAGCTGAAGATGTGT | RT-qPCR, *E* = 97.3% |
| *BgElo23* (110) | ACTGGGCGTGGATGTTCTAC | GCCCGCATGATGATACGACT | RT-qPCR, *E* = 99.8% |
| *BgElo24* (147) | TGGCTGGTCTCTTGGATTGG | ATCGGGGTCCCACAACATTC | RT-qPCR, *E* = 96.9% |
| *BgDsx* (113) | TGCACAGGCGCAGGACGAGG | CCTCCATGATGGCTGCGGTG | RT-qPCR, *E* = 101.1% |
| *BgTra* (83) | CCACCCATGAGAGAAAGGGA | GGACCATCTGGTGATTTCGATCT | RT-qPCR, *E* = 101.6% |
| *Actin5C* (95) | GCTTTGCTATGTTGCCCTCG | CCTGACCATCAGGAAGCTCG | RT-qPCR, *E* =102.0% |
| *BgElo1* (434) | T_7_ + GGGATGGGCAACACGTTATC | T_7_ + GGTAGCCACAATCCTTCGTG | dsRNA synthesis |
| *BgElo2* (343) | T_7_ + GGCCCAGGATGATGGAAAAC | T_7_ + GGAGCAAACTTCATGCCCAA | dsRNA synthesis |
| *BgElo3* (420) | T_7_ + AGAGCGATCCTCGTGTGAAT | T_7_ + ACCAGACACTCATGGGCATA | dsRNA synthesis |
| *BgElo6* (293) | T_7_ + TGCCAACCAACACAAAGGG | T_7_ + TTTGGTCCCATTGCTGCTAC | dsRNA synthesis |
| *BgElo7* (344) | T_7_ + ATGATGGCTGCTATGGGACC | T_7_ + TCATCAGTGCAAGGAGACAGA | dsRNA synthesis |
| *BgElo9* (419) | T_7_ + ATGGAGATCCACGCACTTCA | T_7_ + GACACTCATCGGCATCACAC | dsRNA synthesis |
| *BgElo10* (349) | T_7_ + GCAGCTTCTTACACATGGCT | T_7_ + TGAAAACCAGGGTACCTGTCA | dsRNA synthesis |
| *BgElo11* (496) | T_7_ + AGGATCACCATGGCCTACAA | T_7_ + TGGCCACATGGAGGTTACAA | dsRNA synthesis |
| *BgElo12* (285) | T_7_ + ACCAAGTCTCGTTTCTTCACG | T_7_ + TTTTGGGCATAGCACAGCTC | dsRNA synthesis |
| *BgElo12B* (361) | T_7_ + GGGAACATCACAGCTCTCAGTA | T_7_ + CCATCTGTCCATCCAGATGTTA | dsRNA synthesis |
| *BgElo14* (377) | T_7_ + TCGGTTGCTTTGTGAACCTG | T_7_ + TGTGCATTGCGATGAGGAAG | dsRNA synthesis |
| *BgElo17* (296) | T_7_ + GGGCTTGGGTCCTCATCTAA | T_7_ + CGTGCAGGAATGTAGCTTGA | dsRNA synthesis |
| *BgElo20* (318) | T_7_ + CAGCTTCGTTGATGGCTACC | T_7_ + TCCGGATAGCTTTGGGGTTT | dsRNA synthesis |
| *BgElo22* (266) | T_7_ + TGGATGGTGAAAAACTGGACA | T_7_ + CAGAACCCTGACACTCGGTC | dsRNA synthesis |
| *BgElo24* (298) | T_7_ + TCAGATAGCCCTTTGCACCT | T_7_ + ACCAACTATGCTCCCATGCC | dsRNA synthesis |
| *BgElo24B* (293) | T_7_ + CTTTGTGACGAATCGGTGGC | T_7_ + AGCAACTTCTCCCATGTGACT | dsRNA synthesis |
| *BgDsx-A* (270) | T_7_ + GCATGTGCGGGAGTGAGAGA | T_7_ + CGTCGTAGCGCCGTCTGTAG | dsRNA synthesis |
| *BgDsx-B* (340) | T_7_ + GTTGCGACTCTTCGTCCTCA | T_7_ + ACGTAGATGAGCGGAAGTGC | dsRNA synthesis |
| *BgTra-A* (207) | T_7_ + AGATTTCATTCAAGTGGACCTGGATCATAC | T_7_ + TCACATACCTGGTCTCCGCT | dsRNA synthesis |
| *BgTra-B* (361) | T_7_ + CATGGGTTCCCGCCTTAGTAAT | T_7_ + TCACATACCTGGTCTCCGCT | dsRNA synthesis |
| *Muslta* (193) | T_7_ + CACCCTCTCCACGAATTG | T_7_ + TAGAAGATGCTGCTGTTTCA | dsRNA synthesis |
| *BgElo12* (825) | CGGGGTACC***TACACA***ATGGCGGCATTAACGACAAG | CGCGGATCCTTATTGCTTATGAGATTTTC | Yeast expression |
| *BgElo24* (958) | CGGGGTACC***TACACA***ATGGCTGGTCTCTTGGATTG | CGCGGATCCTGACCGCTTTTGAACGCAAT | Yeast expression |
| *GFP* (729) | CGCGGATCC**TACACA**ATGGTGAGCAAGGGCGAGGAG | CCGGAATTCTTATTTGTATAGTTCATCCA | Yeast expression |
| *BgDsx1* (1095) | CGGGGTACCATGTCGGAGAACGGCAGCGAGAC | CCGCTCGAGCTAGTGGTGGTGGTGGTGGTGCGAAGCTGGTGGTTCGCTGCTAGG | Dual-luciferase reporter assay |
| *BgTra5* (2061) | CCGGAATTCATGCGATCAAGGTCTCCAAGTCGC | CCGCTCGAGCTAGTGGTGGTGGTGGTGGTGCTTGTCAACATCATTGTTGCTAAGC | Dual-luciferase reporter assay |
| *BgTra6* (1827) | CCGGAATTCATGCGATCAAGGTCTCCAAGTCGC | CCGCTCGAGCTAGTGGTGGTGGTGGTGGTGCTTGTCAACATCATTGTTGCTAAGC | Dual-luciferase reporter assay |
| *BgElo12*-GSP |  | GATTACGCCAAGCTTGTACCGGAGGAGAGCATCCAAGCGT | 5’ RACE |
| *BgElo12* (2678) | CGGGGTACCAGGTTCAAACACGGGGCTGACC | CCGCTCGAGACTGAGAGCTGTGATGTTCCCTTCTC | Dual-luciferase reporter assay |

T_7_ means the T_7_ promoter sequence: GATCACTAATACGACTCACTATAGGG.

The amplification efficiency of qPCR is represented by *E*.

The underlined sequences in yeast expression primers represent the restriction cleavage sites, the sequence in bold italics represents the yeast consensus sequence.

The underlined sequences in dual-luciferase reporter assay primers represent the restriction cleavage sites, green sequences represent termination codon, red sequences represent His-tag.
